# Simplified model assumptions artificially constrain the parameter range in which selection at the holobiont level can occur

**DOI:** 10.1101/2020.03.14.988766

**Authors:** Itay Daybog, Oren Kolodny

## Abstract

Van Vliet & Doebeli (1) (henceforth VVD) present a multi-level selection framework simulating host-microbiome evolutionary dynamics. The model explores the conditions under which the association between hosts and their “helper” microbiome – microbes that developed a trait that provides a benefit to the host at a cost to themselves – is strong enough to allow for selection at the holobiont level. The model is an important contribution to the holobiont debate, making the notion quantifiable and highlighting the involved parameters.

In the study, VVD conclude that the parameter space in which selection occurs at the level of the holobiont is extremely restricted (Fig 1). Here we show that this result stems from specific model assumptions. Although these assumptions are reasonable as a modeling starting point, slight biologically-reasonable modifications of the assumptions lead to qualitatively different outcomes.

## Main Text

One model-assumption that can readily be altered is the relation between microbiome composition and host fitness. In VVD’s model, the probability to reproduce increases linearly with the quantity of the helper microbes. Many alternatives are reasonable, reflecting different inter-generational selection dynamics of the host.

One plausible scenario is that the fitness acts as a step function. This may occur, for example, if the helper microbiome supplies a vital nutrient, otherwise inaccessible to the host, that is required in small amounts (2). To test this, we ran VVD’s model with host fitness determined by a discrete step function, at a threshold of 1% helper microbes. Individuals whose helper frequency in the microbiome was above the threshold received fitness 0.95, and others 0.05. This simple alteration of VVD’s model-assumptions leads to a broad parameter space in which a stable population of helper microbes is maintained despite their costly behavior, suggesting selection at the holobiont-level (Fig 2). Such a non-linear dependency of fitness on an underlying factor that can be influenced by the microbiome is not uncommon in the real world, stemming from physiological, ecological, or behavioral factors (2–5). VVD’s model may thus suggest the opposite conclusion from the one they reach, depending on the choice of fitness function.

**Fig. 1.**
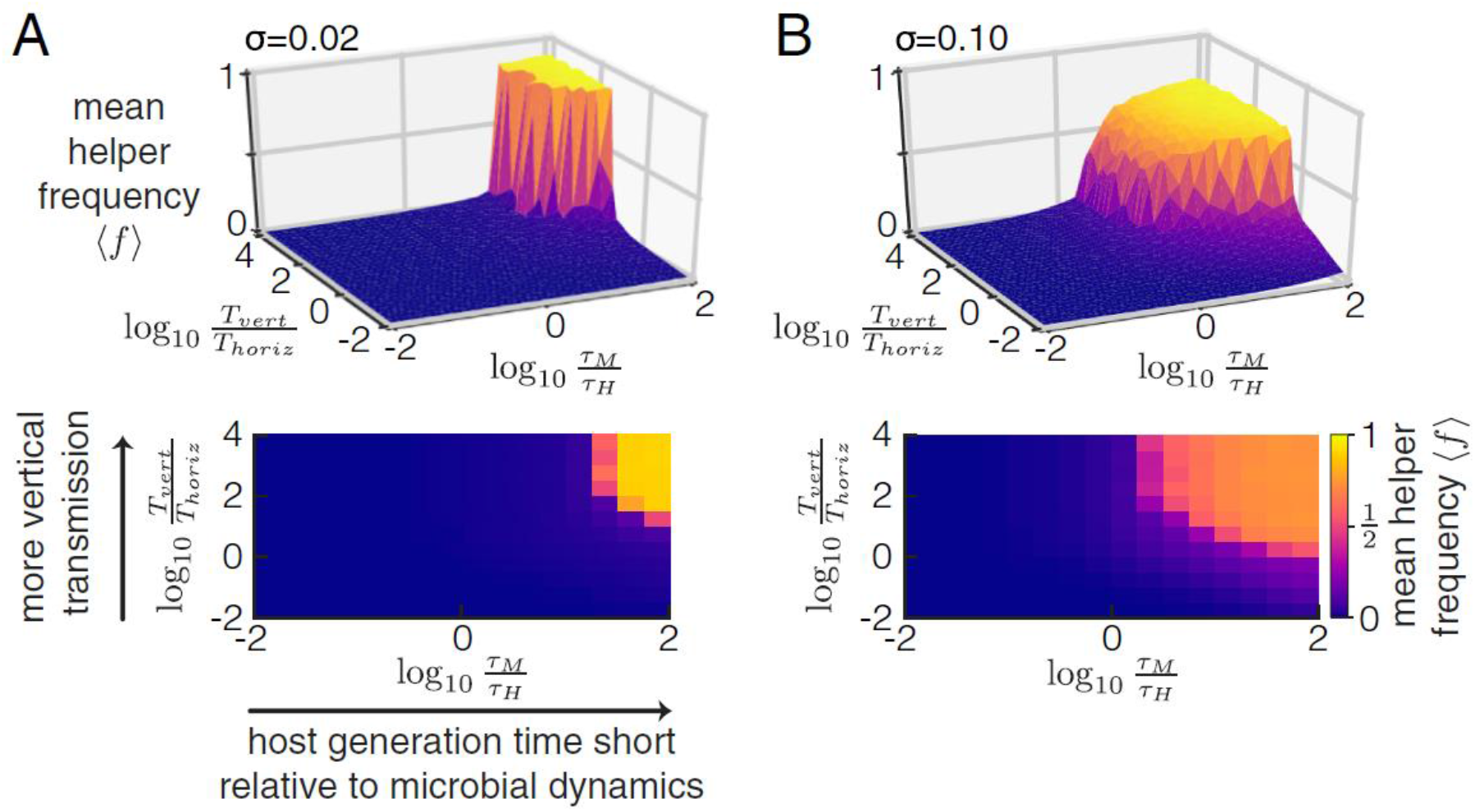
VVD’s original results - Selection can only act at the holobiont level under stringent conditions. The mean helper frequency is shown as a function of the timescale of the evolutionary dynamics at the microbe level relative to the host generation time and the ratio of vertical to horizontal transmission. Two different levels of sampling variance are shown: the SD of the sampling distribution is 0.02 in A and 0.1 in B; A (top) and B (top) show the results of single simulations, and A (bottom) and B (bottom) show the same data, averaged within discrete bins.

**Fig. 2.**
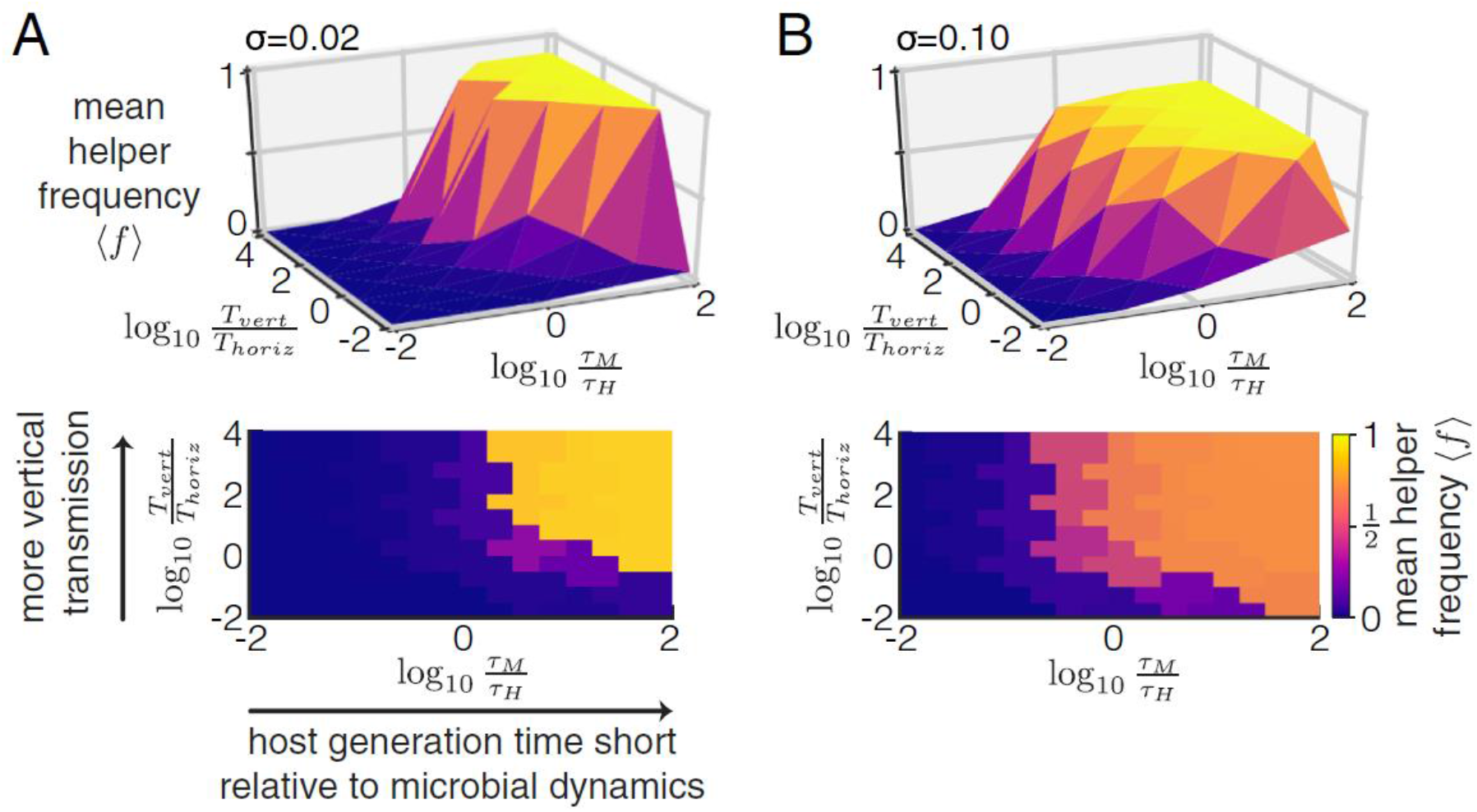
When replacing the assumed fitness function by a simple, biologically-reasonable, step function, VVD’s model predicts that selection would act at the level of the holobiont across a much broader range of parameter values than in VVD’s original findings (compare each panel to its counterpart in figure 1; panel descriptions analogous to those of Figure 1).

Other assumptions of VVD’s model may be altered to reflect realistic dynamics. VVD discuss the possibility that the host controls its microbiome composition to some extent. However, they do not model such a possibility due to its complexity. This is reasonable, but respectively warrants caution when drawing conclusions about reality from the model. We suggest that related general scenarios can be easily integrated into the model. For example, a host may be able to flush out its microbiome if its helper frequency is too low, allowing re-establishment with a random draw of new microbes (6). This is likely to have a dramatic effect on the model outcomes. Another aspect implicitly assumed in VVD’s model is its combination of hard and soft selection; modeling selection as primarily soft – a common situation in reality – is also likely to alter the model’s results.

The framework proposed by VVD provides a useful demonstration of the way to constructively tackle questions of host-microbiome evolutionary dynamics with a simplified model that captures essential components of the system. In our view, the assumptions that were chosen in their implementation are an appropriate choice for an initial exploration. However, they are far too limiting and simplistic to derive general conclusions about the likelihood of selection at the holobiont level, warranting cautious interpretation and calling for further exploration.

Scripts, numerical results, and details are included in documentation available at https://github.com/itaydaybog/holobiont-multilevel-selection-response. Supplementary code can be found below. We thank the Kolodny lab members for insightful comments, and the Gordon and Betty Moore Foundation (Symbiosis in Aquatic Systems initiative), for funding.

## Supplementary code and files analysis

### Relevant code snippets

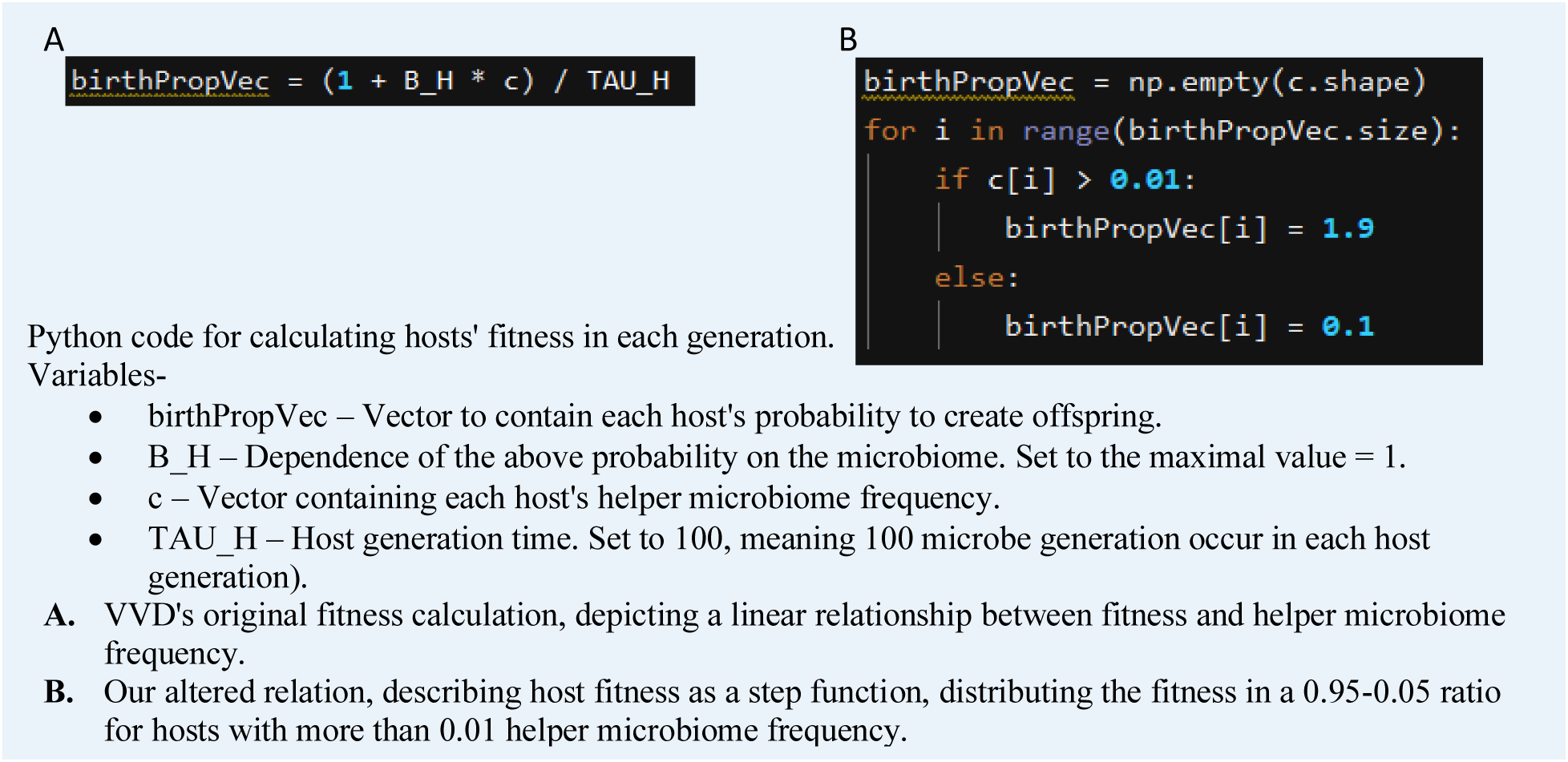

### Git files

In the repository https://github.com/itaydaybog/holobiont-multilevel-selection-response are the following:

- Model code

- VVD’s original files

- mls_general_code_original.py - Helper functions used by the model.
- MLS_static_fast_original.py - Runs the original model.
- Our files

- MLS_static_fast_new.py - Built on VVD’s original model but implements the fitness as the step function.
- Figure generating code

- VVD’s original files

- MLS_figure_3_original.py - Generates figure 3 in VVD’s manuscript.
- Our files

- MLS_figure_3_new_single_point.py - Runs our altered model on a single point in the parameter space.
- build_3d_graph.py - Uses all the data created by MLS_figure_3_new_single_point.py to build a continuous representation of the results.
- Data files

- Original_Data - Data outputted by VVD’s original model, by running “MLS_figure_3_original.py”.
- Step_Data - Data outputted by our edited model, by running “MLS_figure_3_new_single_point.py” on several points in the parameter space.
- Step_figures & Original_Figures - Empty folders to contain figures created by the models.

